# Spontaneous Eye Blink Rate during the Working Memory Delay Period Predicts Task Accuracy

**DOI:** 10.1101/2021.03.15.435433

**Authors:** Jefferson Ortega, Chelsea Reichert Plaska, Bernard A. Gomes, Timothy M. Ellmore

## Abstract

Spontaneous eye blink rate (sEBR) has been linked to attention and memory, specifically working memory (WM). sEBR is also related to striatal dopamine (DA) activity with schizophrenia and Parkinson’s disease showing increases and decreases respectively in sEBR. A weakness of past studies of sEBR and WM is that correlations have been reported using blink rates taken at baseline either before or after performance of the tasks used to assess WM. The goal of the present study was to understand how fluctuations in sEBR during different phases of a visual WM task predict task accuracy. In two experiments, with recordings of sEBR collected inside and outside of a magnetic resonance imaging bore, we observed sEBR to be positively correlated with WM task accuracy during the WM delay period. We also found task-related modulation of sEBR, including higher sEBR during the delay period compared to rest, and lower sEBR during task phases (e.g., stimulus encoding) that place demands on visual attention. These results provide further evidence that sEBR could be an important predictor of WM task performance with the changes during the delay period suggesting a role in WM maintenance. The relationship of sEBR to DA activity and WM maintenance is discussed.

## 1 Introduction

The healthy human blinks around 15-20 times per minute (Tsubota et al., 1996), however the precorneal tear film, which lubricates the eye, only begins drying up approximately 25s after a blink ends (Norn, 1969). This suggests that we blink more often than needed to maintain a lubricated precorneal tear film. Previous research has found task-related modulation of spontaneous eye blink rate (sEBR), which indicates that blinking could be reflective of cognitive factors (Siegle et al., 2008;Oh et al., 2012). For example, reading is accompanied by low levels of sEBR, while high levels of sEBR have been reported during conversation (Bentivoglio et al., 1997). More recent studies have found that sEBR correlates with attentional load and fatigue (Maffei and Angrilli, 2018), attentional control (Colzato et al., 2009;Unsworth et al., 2019a), can track working memory updating and gating (Rac-Lubashevsky et al., 2017), and can predict differences in exploration during reinforcement learning (Van Slooten et al., 2019). In addition, a growing body of literature continues to provide evidence supporting sEBR as an effective measure of striatal dopamine (DA) activity (Jongkees and Colzato, 2016). However, whether sEBR does indeed reflect DA activity is still debated today (Dang et al., 2017;Sescousse et al., 2018).

The connection between sEBR and DA first came from observations of neurological and psychiatric disorders that found decreased sEBR in patients with Parkinson’s (Hall, 1945;Reddy et al., 2013), a neurodegenerative disorder that affects the dopaminergic system in the brain, causing symptoms like rigidity (Dauer and Przedborski, 2003). Schizophrenia has also been suggested to provide evidence for a connection between sEBR and DA due to excessive DA activity in the striatum (Howes et al., 2015) and increased sEBR in schizophrenia patients (Adamson, 1995;Swarztrauber and Fujikawa, 1998). Additionally, sEBR and DA has previously been investigated in pharmacological studies, which have observed an increase in sEBR after administration of DA agonists, while DA antagonists decreased sEBR in primates (Elsworth et al., 1991;Jutkiewicz and Bergman, 2004). In one study, researchers found sEBR was correlated with dopamine levels specifically in the caudate nucleus in monkeys, suggesting that DA could regulate blink rate (Taylor et al., 1999). This is further supported by another study that found sEBR to be closely related to *in vivo* and positron emission tomography (PET) measures of striatal D2 receptor density in the ventral striatum and caudate nucleus of adult male vervet monkeys (Groman et al., 2014). These findings provide valuable evidence for sEBR being a viable measure of striatal DA activity and has led to many researchers to adopt sEBR as a measure of DA activity. Moreover, sEBR is an easy-to-record physiological measure that is non-invasive and affordable.

One cognitive process of interest, that is also closely related to DA activity, is working memory (WM) which is the process of actively holding information online and manipulating it to meet task demands (Baddeley, 1992). Prior research has found substantial evidence that demonstrates the importance of dopaminergic neurotransmission and the role of the prefrontal cortex during WM function (Fuster and Alexander, 1971;Funahashi et al., 1989;Courtney et al., 1998;Wager and Smith, 2003;Cools and Robbins, 2004), especially during WM maintenance (Fuster and Alexander, 1971;Funahashi et al., 1989;Constantinidis et al., 2018). Specifically, human studies investigating DA in WM tasks have found both caudate dopamine activity during WM maintenance and DA synthesis capacity to be positively correlated with WM capacity, a measure of the amount of information that can be held in WM (Cools et al., 2008;Landau et al., 2009). Though it is widely accepted that the PFC plays an important role in WM function (Roberts et al., 1998), many researchers still debate the PFC’s role in WM (Seamans and Yang, 2004). One model that attempts to elucidate the PFC’s role in WM function is the prefrontal cortex basal ganglia WM model (PBWM) (Frank et al., 2001;Hazy et al., 2006). PBWN is a computational neural network model that suggests that WM requires robust maintenance and rapid selective updating. This model states that the frontal cortex facilitates robust, active maintenance through recurrent excitation in frontal neurons, while the basal ganglia orchestrates a gating mechanism that controls the flow of information into WM (Frank et al., 2001). Previous research has pointed towards DA being important for this sustained firing activity in the PFC during WM maintenance (Sawaguchi, 2001;Durstewitz and Seamans, 2008;de Frias et al., 2010). The relationship between DA and WM performance is believed to follow an inverted-U-shape, in which too little or too much dopamine impairs performance, as seen in psychopharmacological studies (Stewart and Plenz, 2006). In one study, the effects of administered dopaminergic drugs on PFC function depended on baseline levels of performance, whereas administration of bromocriptine, a dopamine agonist, impaired performance for individuals with higher working memory abilities while improving performance for individuals with lower working memory abilities (Kimberg et al., 1997).

Although sEBR has been used in prior research to investigate cognitive functions like WM, many of these studies relied on baseline levels of sEBR to investigate these relationships (Tharp and Pickering, 2011;Zhang et al., 2015;Unsworth et al., 2019b). Few studies have investigated the relationship between phasic sEBR during a WM task. Phasic sEBR refers to the measuring of sEBR in response to stimulus conditions while tonic sEBR refers to baseline levels of blinking (Bacher and Allen, 2009). To the best of our knowledge, only one other study has examined sEBR as a function of the different task phases (e.g., stimulus encoding, maintenance during the delay period, and stimulus probe periods) of a WM task (Bacher et al., 2017). Bacher et al. (2017) found modulation of sEBR across these different phases are developed in infants as young as 10 months, indicating that sEBR can reflect dopamine function in early human development. They also observed higher sEBR during the Hide (delay) phase of the task in relation to the Reveal phase, which is when the experimenter revealed the toy’s location to the child. This modulation of sEBR was suggested to reflect the engagement of cognitive resources that have become available during the Hide phase and transiently elevated DA activity that is needed to update and maintain mental representations (Bacher et al., 2017).

The goal of the current study was to investigate how fluctuations in sEBR during different phases of a Sternberg visual WM task (Figure 1) relate to performance, and how sEBR fluctuations change across task demands. First, we hypothesized that sEBR during the WM Delay period, when stimuli are being maintained, would be positively correlated with task performance and that there would be a non-linear relationship such that low and high sEBR would correlate with worse performance. Second, we hypothesized differences in sEBR across phases of the WM task with differences between phasic sEBR during the WM delay and tonic sEBR during non-task rest periods.

**Figure 1.**
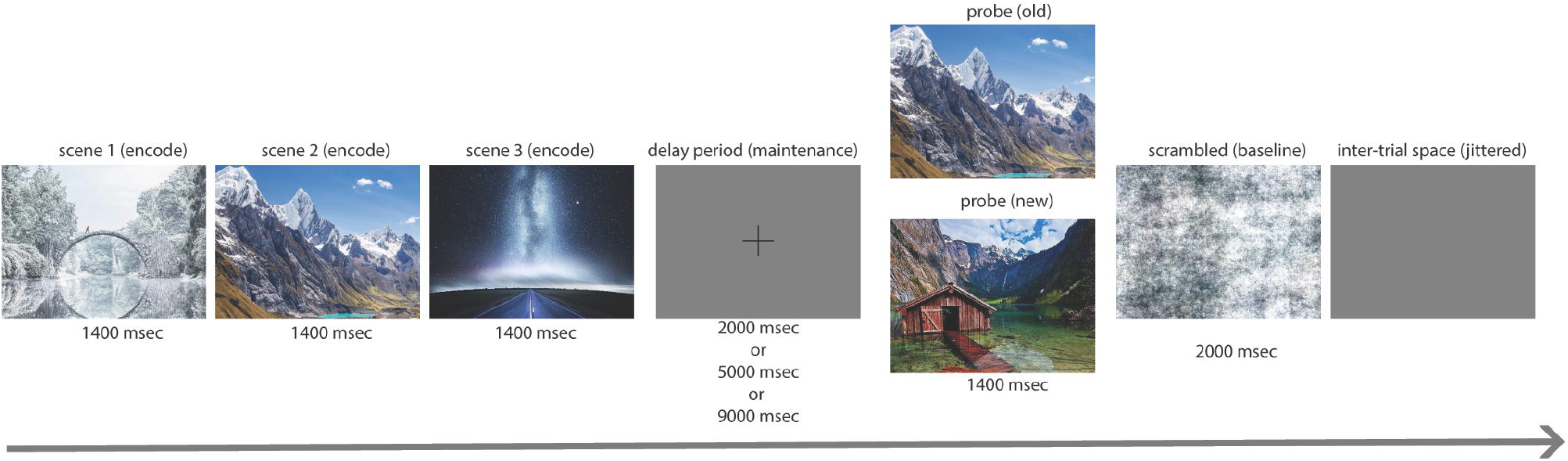
Task design. Each trial began with an encoding period where three novel complex scenes were presented for 1400 ms each. The encoding period was followed by a delay period where a fixation cross was presented on a grey background for a varied amount of time (2 s, 5 s, or 9 s). After the delay period, the probe was presented for 1400 ms and participants had to identify whether the image was a new image or one of the previously presented images with a button press. After the probe, a scrambled image was presented for 2000 ms which indicated the end of the trial followed by jittered blank space before the start of the next trial.

## 2 Materials and Methods

### 2.1 Participants

The experiments were conducted under a protocol approved by the Institutional Review Board of The City University of New York Human Research Protection Program (CUNY HRPP IRB). All methods were carried out in accordance with the relevant guidelines and regulations of the CUNY HRPP IRB committee. All participants were recruited either by flyers posted throughout the City College of New York campus or by web postings on the City College of New York SONA online experimental scheduling system. All participants had normal or corrected-to-normal vision with no reported neurological or psychiatric disorders. Participants were either compensated $15 per hour or received one psychology course credit per hour of participation in the study. Written informed consent was obtained from all participants in the study.

Participants selected for Experiment 1 and Experiment 2 were part of a larger study. Nineteen healthy participants (8 males; M = 23.79; S.D. = 7.72) were recruited for Experiment 1. In Experiment 1, sEBR was measured inside a 3 tesla Siemens Prisma MRI scanner. In Experiment 2, sEBR was recorded in a sound attenuated EEG booth during acquisition of EEG data while participants sat upright. Fifty-three healthy participants (29 males; M = 23.58; S.D. = 5.79) were recruited for Experiment 2. Three participants were removed from Experiment 1 including one participant who was removed for noisy data and two who were removed for task performance below or close to chance. A total of 19 participants were removed from Experiment 2 for multiple reasons including 11 participants who were removed due to bad EOG channel quality, four participants who were removed because of a stimulus marker malfunction, three participants who were removed due to outlier detection, and two who were removed for failing to adhere to the protocol. The final sample for the analysis in Experiment 1 was sixteen subjects, and for Experiment 2 was thirty-four subjects.

### 2.2 Task and Procedure

Prior to the start of the task, participants completed a 5-minute Rest period which consisted of staring at a black fixation cross that was shown on a grey background. Participants completed another 5-minute Rest period after completing three runs of the task. This fixation cross was also used during the delay period of the task. Participants completed three runs, each run containing 54 trials, of a modified version of the Sternberg WM task (Sternberg, 1966). Naturalistic scenes were used as stimuli and were sampled from the SUN database (Xiao et al., 2010). The task consisted of a stimulus encoding period, delay period, probe period, and post-probe scrambled stimulus period (which served as a visual baseline and to signal end of trial). During the encoding period, participants were shown three subsequent novel scenes for 1400 ms each. During the delay period, a black fixation cross was shown on a grey background for varied lengths (either 2, 5, or 9 seconds long). The delay period duration was randomized from trial to trial to engage subjects’ attention consistently across trials because they could not predict when the delay period would end. Each three runs of the task had 18 trials of each delay duration with order of presentation randomized. The probe was presented for 1400 ms after the delay period and consisted of a new image (one that has not been presented yet) or an old image (one that was shown during encoding). The chance of receiving a new probe was 50%. Participants indicated whether the image presented was either a new or an old image with a button press. After the probe, a Fourier phase-scrambled scene was shown for 2000 ms, indicating the end of the current trial followed by a jittered period of blank screen.

### 2.3 sEBR Recording

Participants were not given instructions about when to blink during the experiment. Previous studies have found blink rate to be stable between 10 am and 5 pm (Barbato et al., 2000;Doughty and Naase, 2006). For both Experiment 1 and Experiment 2, sEBR was recorded between 10:00 am and 3:00 pm. During Experiment 1, eye blinks were recorded inside a 3 tesla Siemens Prisma MRI scanner using an MRI compatible EyeLink 1000 Eye Tracker (SR Research) and was recorded at 500 Hz. In Experiment 2, eye blinks were recorded using an electrooculogram (EOG) that was recorded during 64-channel scalp electroencephalogram (EEG) using a Brain Products cap and active electrode recording system (actiChamp). EOGs were placed above the left eye and below the right eye to track blinking. Blink detection was performed using MNE Python via the function “*find_eog_events*” (Gramfort et al., 2013). Blink epochs were evaluated for each run of the task for all participants. Runs with blink epochs which did not resemble the standard blink shape were removed from the analysis. Only participants with 2 or more runs of good eye-tracking data were used in the analysis. The first two seconds of all delay periods were used in the analysis. In Experiment 1 and 2, sEBR was computed by dividing the total number of blinks by the total period duration for any given phase resulting in units of blinks per minute:

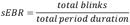

### 2.4 Statistical analysis

Statistical analyses were computed using JASP Version 0.14.1. sEBR and task accuracy data were checked for outliers prior to analysis and were removed. Because the relationship between sEBR and WM performance is believed to follow an ‘inverted U-shape’, we did not consider Pearson’s r the optimal measure for this analysis because it is limited to evaluating only a linear relationship between two variables. Instead, we computed Spearman’s rho, which can describe monotonic functions where as the value of one variable changes the other variable changes but not necessarily at a constant rate. We also used polynomial regression analysis, which is more appropriate for quantifying non-linear associations. Specifically, we investigated the non-linear relationship between task accuracy and sEBR during the Delay period of the task for both Experiment 1 and Experiment 2 using a 2^nd^ order polynomial regression model. Post-hoc statistical power calculations were computed for each experiment and the combined samples of both experiments with G*Power Version 3.1.9.6 using the correlation between sEBR during the Delay period and task accuracy. Parameters included the Exact test family, Correlation: Bivariate normal model with an α error probability of 0.05, the sample size, and the correlation coefficient as the effect size.

## 3 Results

### 3.1 Experiment 1

In Experiment 1, we examined the relationship between sEBR and WM task performance while the duration of the WM delay period interval was varied. The first two seconds of all Delay periods were used in the analysis. First, because of the previously reported non-linear relationship between DA and WM task performance (Cools and D’Esposito, 2011), Spearman’s rho correlation coefficient (*r*_s_) was used to analyze the relationship between sEBR and task accuracy. After performing Bonferroni multiple comparisons correction on p-values, we found no significant relationship between sEBR during the phases of the task and task accuracy (Figure 2a). However, there was a strong positive correlation between sEBR during the Delay period and task accuracy (*r*_s_ = .526, *p* = .036) (Figure 2a). We then examined the correlation between sEBR during the whole trial period and task accuracy to make sure that this relationship was not driving the relationship between sEBR during the Delay and task accuracy. There was no significant relationship between sEBR during the whole trial and task accuracy (*r*_s_ = .149, *p* = .582) (Figure 2b). Descriptive statistics for Experiment 1 are presented in Table 1. Second, we computed a repeated measures ANOVA test to compare participants’ sEBR across the task phases. A Mauchly’s test of sphericity was first computed to check the assumption of sphericity in the data and was found to be significant (*p* = .012). Greenhouse-Geisser and Huynh-Feldt ε values were smaller than 0.75 so a Greenhouse-Geisser correction was performed. There were significant differences in sEBR between group means (F (1.948, 29.213) = 33.196, *p* < .001). A post hoc test using the Holm correction revealed that sEBR was significantly lower during Encoding (18.9 ± 11.0 sEBR, *p* < .001) and Probe (11.3 ± 8.0 sEBR, *p* < .001) periods compared to the Delay period (39.4 ± 19.5 sEBR) (Figure 3). There was no significant difference in sEBR between the Delay and Scrambled period (*p =* 0.682). sEBR was also significantly lower during Encoding (18.9 ± 11.0 sEBR, *p* < .001) and Probe (11.3 ± 8.0 sEBR, *p* < .001) periods compared to the Scrambled period (40.9 ± 15.2 sEBR) (Figure 3). Finally, we investigated the difference between phasic sEBR during the Delay period and tonic sEBR during the Rest period. We performed a paired samples T-test to compare sEBR during the Delay and during Rest. We observed sEBR to be significantly higher during the Delay period (39.4 ± 19.5 sEBR) compared to the Rest period (28.6 ± 14.7 sEBR), t(15) = 2.885, p = .0011 (Figure 4a). We then investigated the correlation between tonic sEBR during the Rest period and task accuracy. There was no significant correlation between sEBR during the Rest period and task accuracy (*r*_s_ = .259, *p* = .333) (Figure 4b).

**Table 1.**
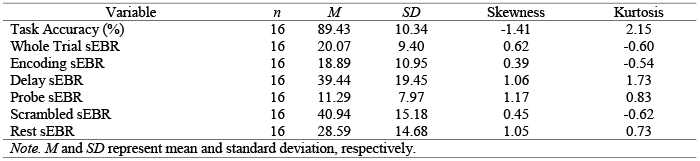
Descriptive Statistics in Experiment 1

**Figure 2.**
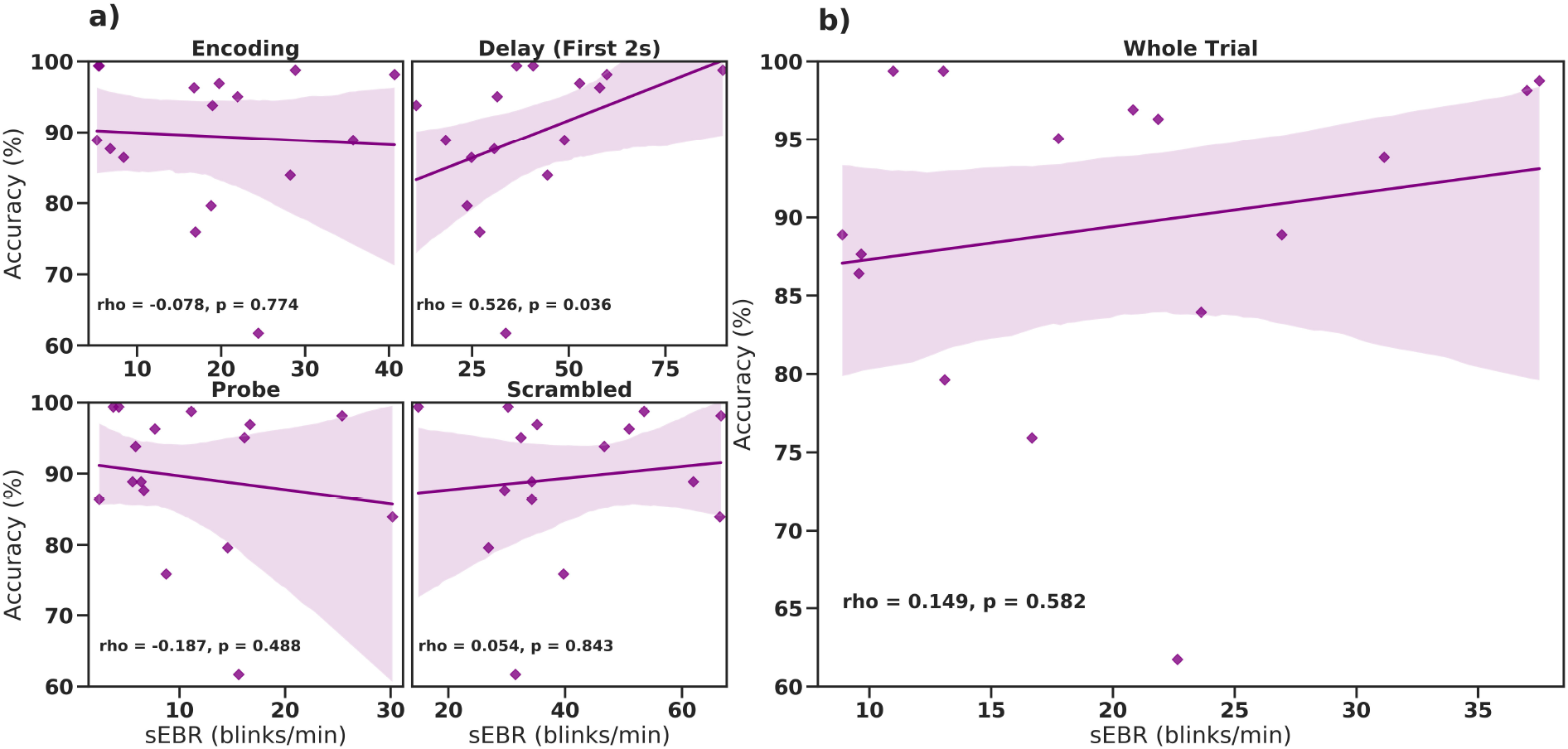
Correlation between sEBR during different phases of the task and task accuracy in Experiment 1. Correlation plots show sEBR (blinks/min) on the x-axis and task accuracy on the y-axis. (**A)** These four plots are encoding (top left), the first two seconds of the delay (top right), probe (bottom left), and scrambled (bottom right) periods. The delay period shows a positive correlation (p=0.036 but not significant after multiple comparisons correction) between task accuracy and sEBR during the first two seconds of the delay period. **(B)** This plot represents the relationship between sEBR during the whole trial and task accuracy. Fitted line represents linear regression model fit. Shaded region depicts 95% confidence interval. **Note:** p-values for Figure 2a. after Bonferroni multiple comparison correction: * p < .0125, ** p < .0025 *** p < .00025. p-values for figure 2b. * p < .05, ** p < .01, *** p < .001.

**Figure 3.**
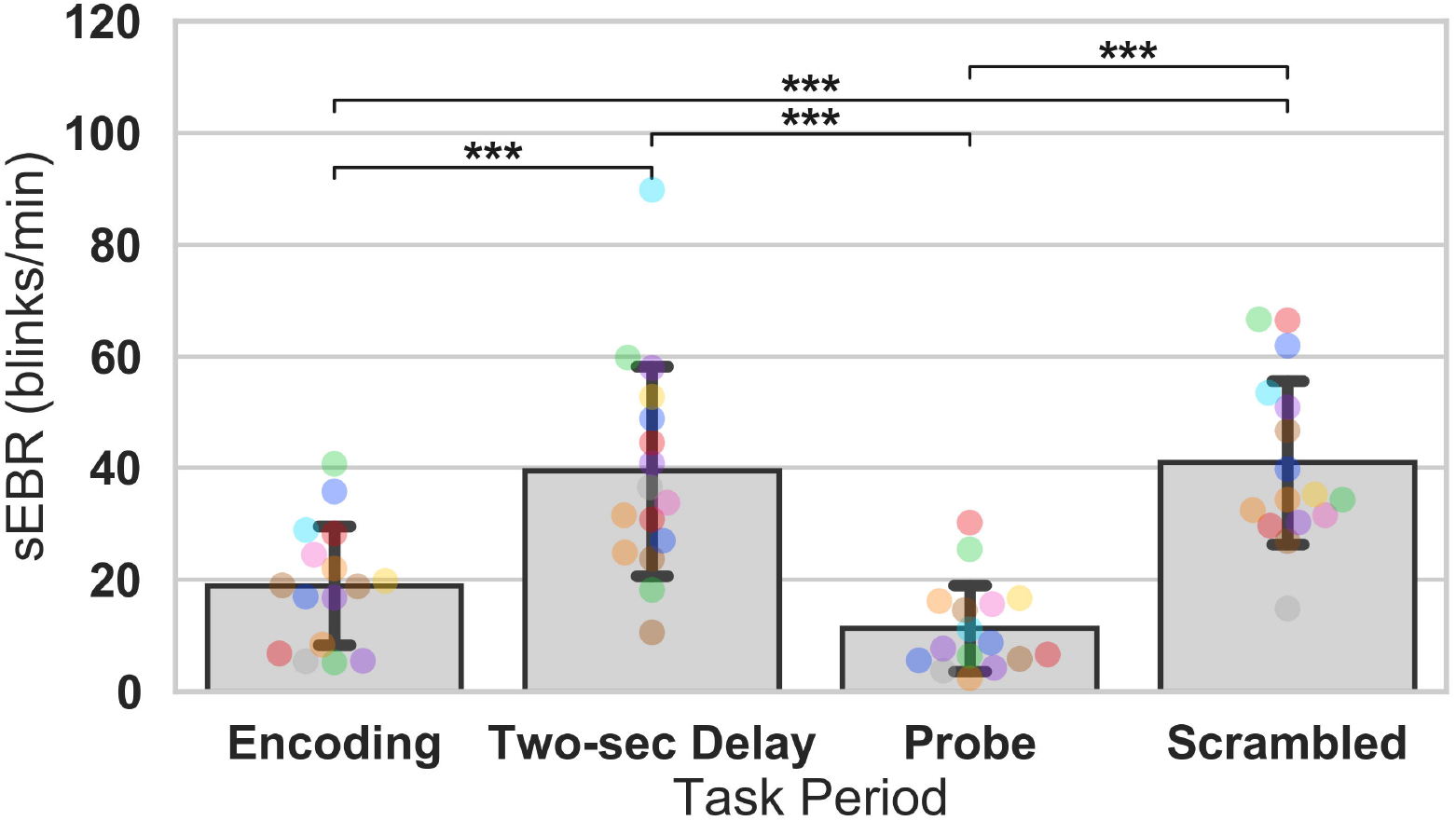
ANOVA test of sEBR across task periods in Experiment 1. Bar plots show task period on the x-axis and sEBR (blinks/min) on the y-axis. Delay period sEBR was significantly greater than Encoding and Probe sEBR. Scrambled sEBR was also significantly greater than Encoding and Probe sEBR. Error bars depict 95% confidence interval. **Note**: Each colored circle represents an individual participant; some colors may be presented twice in one bar due to limited primary colors available for display. P-values were adjusted for comparing a family of 4. * p < .05, ** p < .01, *** p < .001.

**Figure 4.**
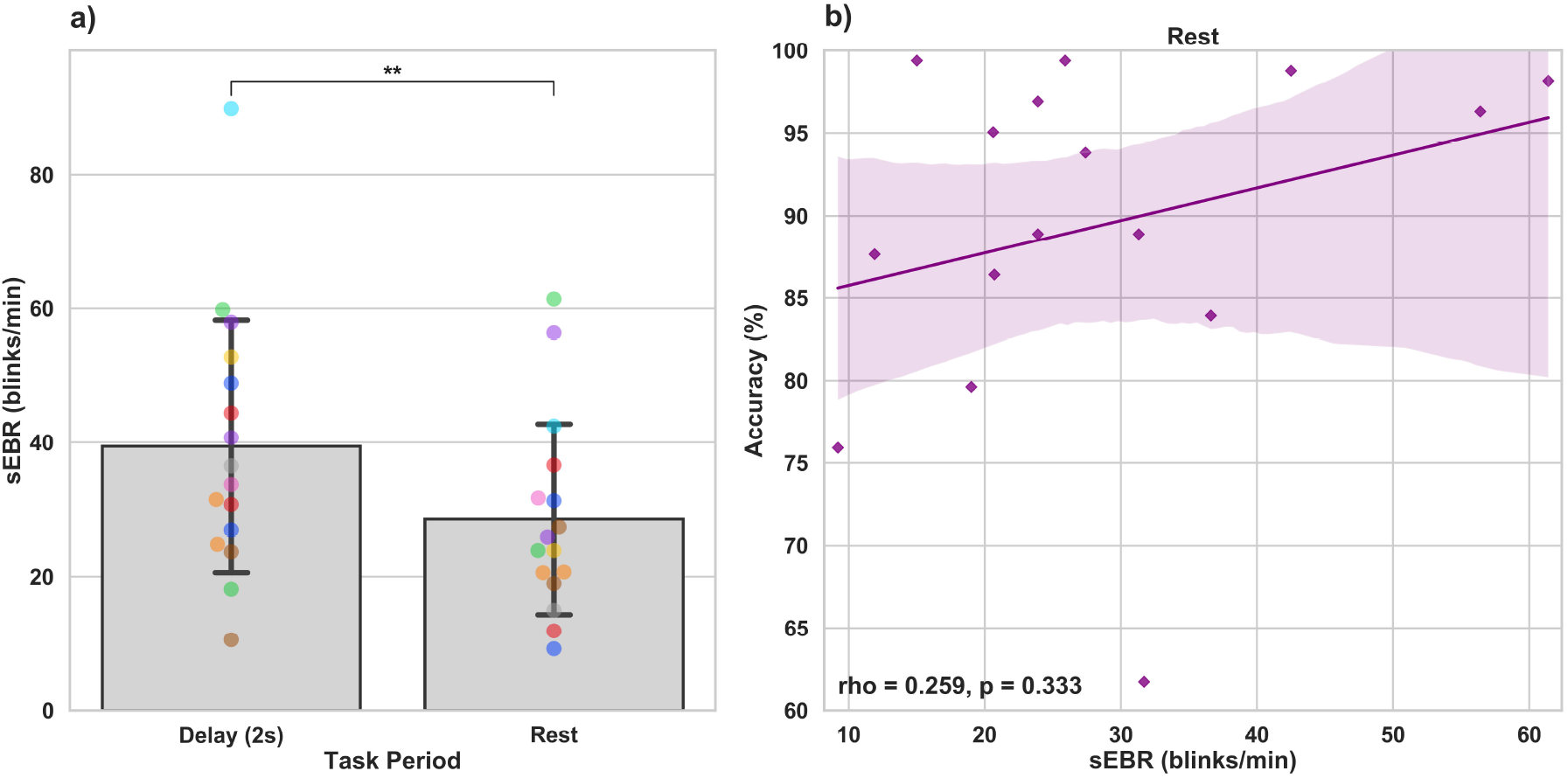
Paired T-tests between Delay period sEBR and Rest sEBR and correlation between Rest sEBR and task accuracy in Experiment 1. **(A)** Bar plots show task period on the x-axis and sEBR on the y-axis. Delay period sEBR was significantly higher than Rest sEBR. Error bars depict 95% confidence interval. (**B)** Correlation plot of sEBR during the Rest period on the x-axis and task accuracy on the y-axis. Fitted line represents linear regression model fit. Shaded region depicts 95% confidence interval. * p < .05, ** p < .01, *** p < .001.

### 3.2 Experiment 2

Experiment 2 included a larger sample of subjects with a task design identical to Experiment 1. First, we examined the relationship between sEBR during each WM task phase and task accuracy. After performing Bonferroni correction on p-values, we found that sEBR during the WM delay period was correlated positively with task performance (*r*_s_ = .508, *p* = .002), with no significant relationships observed between sEBR in other task periods and task performance (Figure 5a). We then examined the relationship between sEBR during the whole trial and task accuracy to make sure that the significant relationship between Delay sEBR and task accuracy was not driven by sEBR during the whole trial. We found no significant relationship between whole trial sEBR and task accuracy (*r*_s_ = .192, *p* = .278) (Figure 5b). Descriptive statistics for Experiment 2 are presented in Table 2. We then repeated the same analysis of comparing sEBR across the task phases by computing a repeated measures ANOVA test. A Mauchly’s test of sphericity was first computed to check the assumption of sphericity in the data and was found to be significant (*p* < .001). Greenhouse-Geisser and the Huynh-Feldt ε values were smaller than 0.75 so a Greenhouse-Geisser correction was performed. There were significant differences in sEBR between group means (F (1.578,52.058) = 66.958, *p* < .001). A post hoc test using the Holm correction revealed that sEBR was significantly lower during Encoding (11.6 ± 8.0 sEBR, *p* < .001), Probe (7.3 ± 4.1 sEBR, *p* < .001), and Scrambled (19.5 ± 10.6 sEBR, *p* < .001) periods compared to the Delay period (35.6 ± 18.3 sEBR) (Figure 6). sEBR was also significantly lower during the Encoding (11.6 ± 8.0 sEBR, *p* < .001) and Probe (7.3 ± 4.1 sEBR, *p* < .001), periods compared to the Scrambled period (19.5 ± 10.6 sEBR) (Figure 6). Additionally, sEBR was significantly lower during the Probe (7.3 ± 4.1 sEBR, *p* = .047), period compared to the Encoding period (11.6 ± 8.0 sEBR) (Figure 6). We then investigated the difference between sEBR during the Delay period and sEBR during the Rest period. We performed a paired samples T-test to compare sEBR during the Delay and during Rest. We observed sEBR to be significantly higher during the Delay period (35.6 ± 18.3 sEBR) compared to the Rest period (17.7± 11.1 sEBR), t(33) = 6.005, p < .001 (Figure 7a). We then investigated the correlation between tonic sEBR during the Rest period and task accuracy. There was no significant correlation between sEBR during the Rest period and task performance (*r*_s_ = -0.053, *p* = .768) (Figure 6b).

**Table 2.**
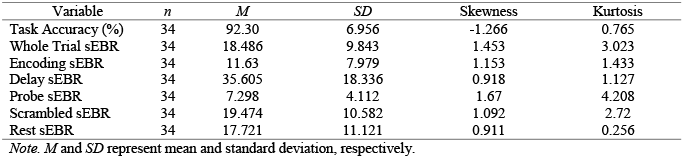
Descriptive Statistics in Experiment 2

**Figure 5.**
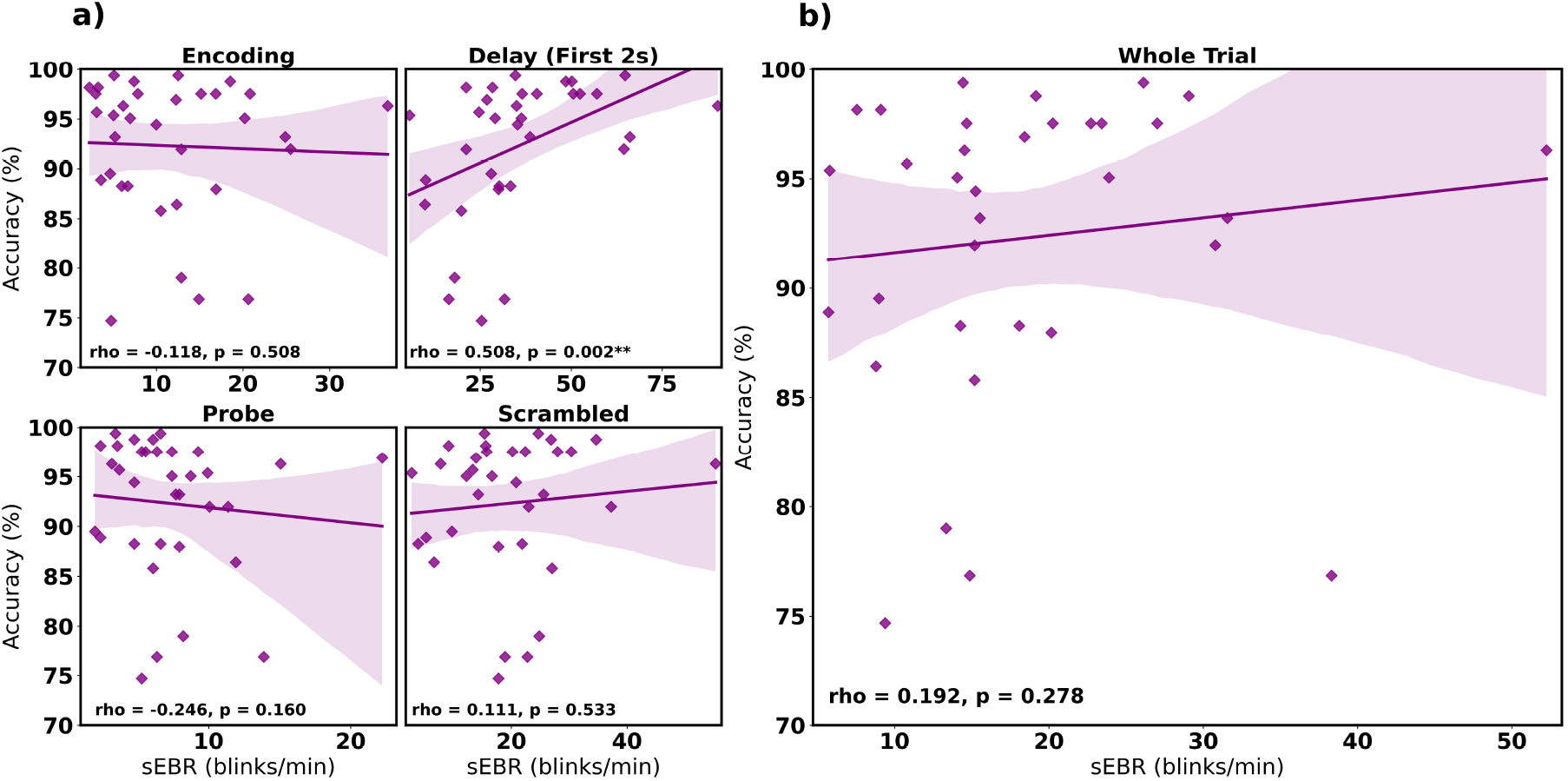
Correlation between sEBR during different phases of the task and task accuracy in Experiment 2. Correlation plots show sEBR on the x-axis and task accuracy on the y-axis. **(A)** These four plots are encoding (top left), the first two seconds of the delay (top right), probe (bottom left), and scrambled (bottom right) periods. The delay period shows a strong positive correlation (p=0.002, significant after a multiple comparison correction) between task accuracy and sEBR during the first two seconds of the delay period. **(B)** This plot represents the relationship between sEBR during the whole trial and task accuracy. Fitted line represents linear regression model fit. Shaded region depicts 95% confidence interval. **Note:** p-values for figure 5a. after Bonferroni correction: * p < .0125, ** p < .0025 *** p < .00025. p-values for figure 5b. * p < .05, ** p < .01, *** p < .001.

**Figure 6.**
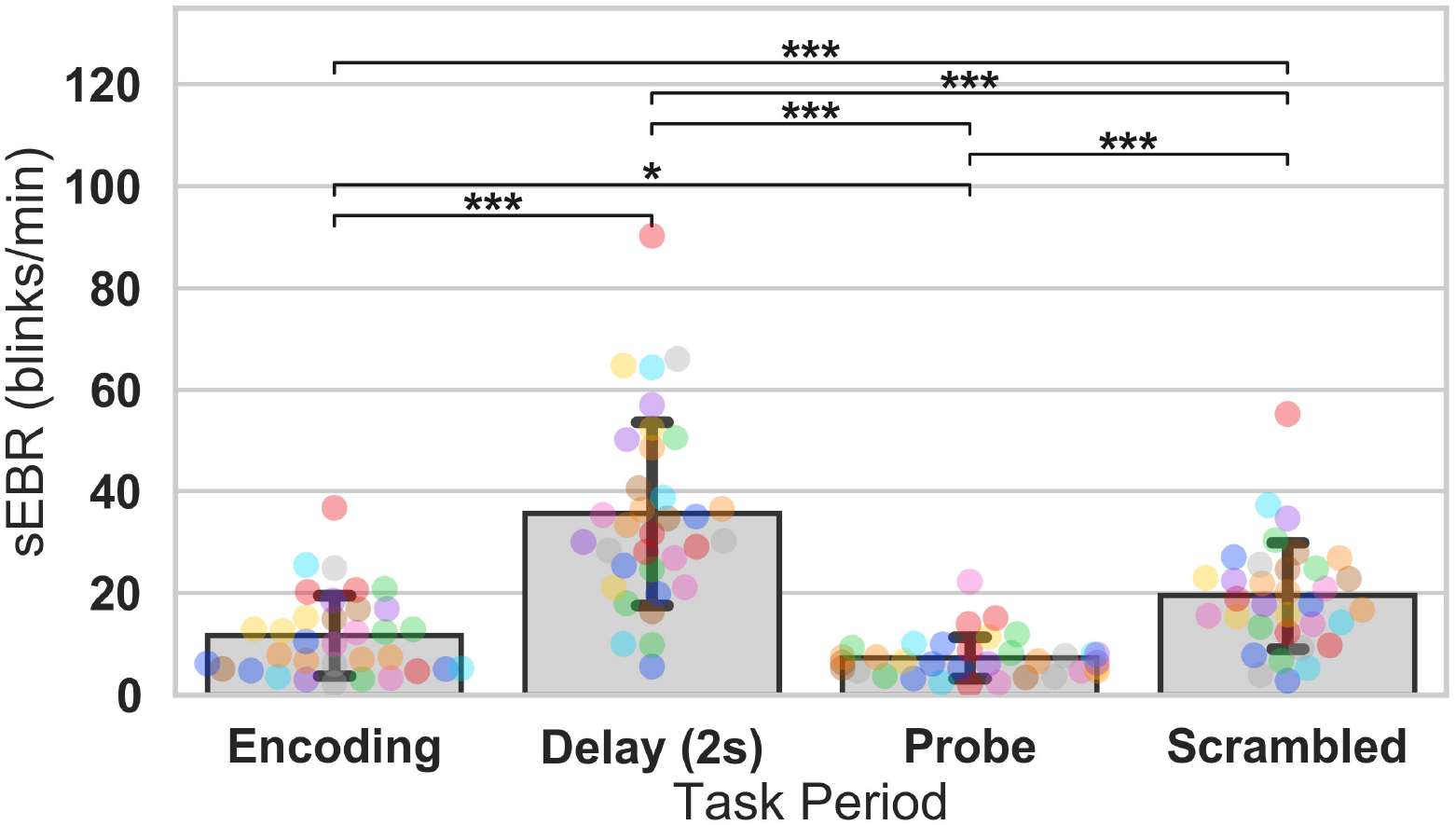
ANOVA test of sEBR across task periods in Experiment 2. Bar plots show task period on the x-axis and sEBR on the y-axis. Delay period sEBR was significantly greater than Encoding, Probe, and Scrambled sEBR. Scrambled sEBR was also significantly greater than Encoding and Probe sEBR. Encoding sEBR was significantly greater than Probe sEBR. Error bars depict 95% confidence interval. **Note**: Each colored circle represents an individual participant; some colors may be presented twice in one bar due to limited primary colors available for display. P-values were adjusted for comparing a family of 4. * p < .05, ** p < .01, *** p < .001.

**Figure 7.**
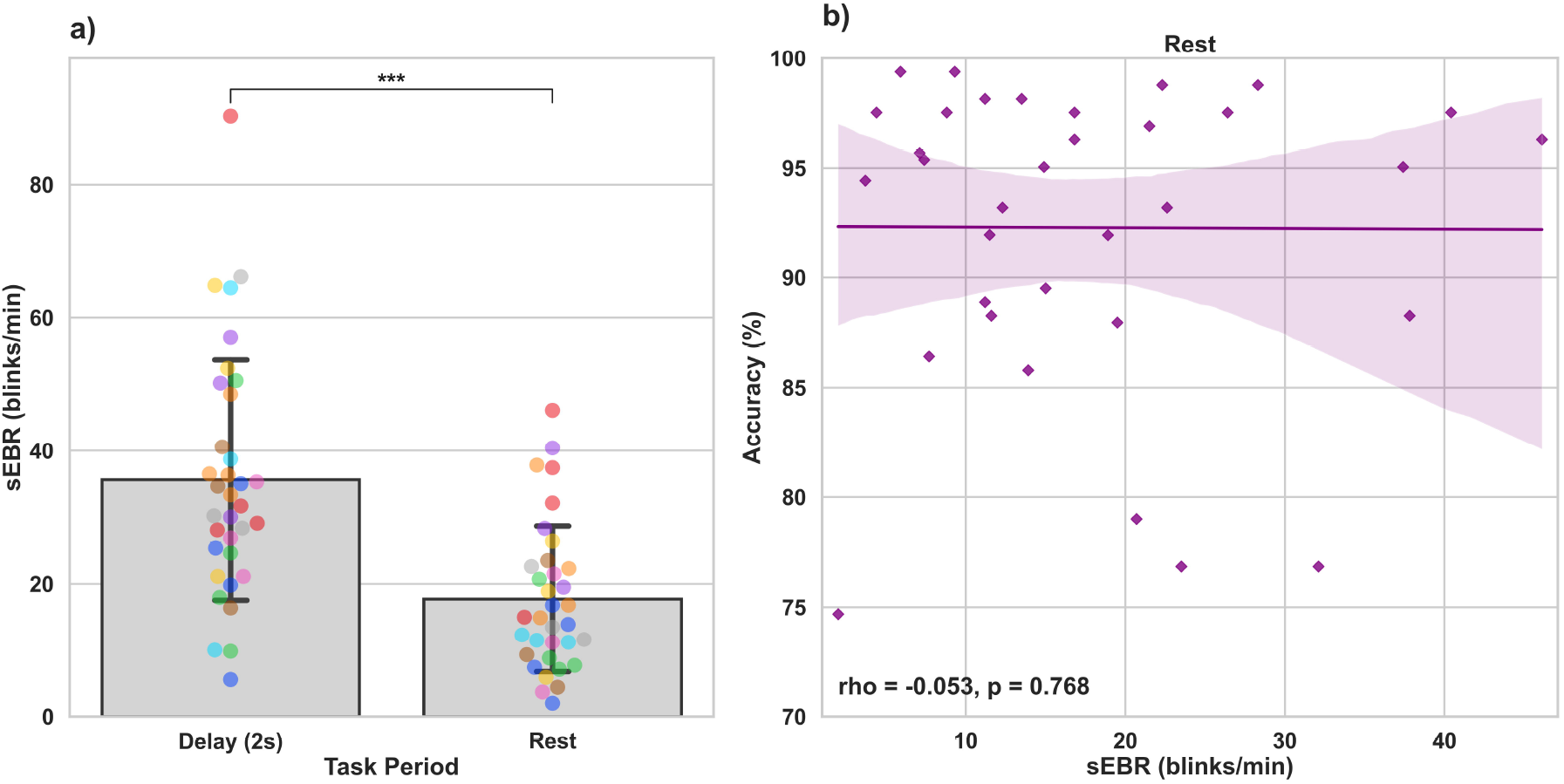
Paired T-tests between Delay period sEBR and Rest sEBR and correlation between Rest sEBR and task accuracy in Experiment 2. **(A)** Bar plots show task period on the x-axis and sEBR on the y-axis. Delay period sEBR was significantly higher than Rest sEBR. Error bars depict 95% confidence interval. **(B)** Correlation plot of sEBR during the Rest period on the x-axis and task accuracy on the y-axis. Fitted line represents linear regression model fit. Shaded region depicts 95% confidence interval. * p < .05, ** p < .01, *** p < .001.

### 3.3 Polynomial Regression Model

To investigate whether sEBR during the Delay varies non-linearly with task performance, we computed a quadratic polynomial regression model between sEBR during the Delay period of Experiment 1 and Experiment 2 and task accuracy. There was no significant polynomial regression relationship found between task accuracy and the first two seconds of the Delay in Experiment 1 (β = 0.324, *p* = 0.741) nor between task accuracy and the first two seconds of the Delay in Experiment 2 (β = -0.568, *p* = 0.322) (Figure 8).

**Figure 8.**
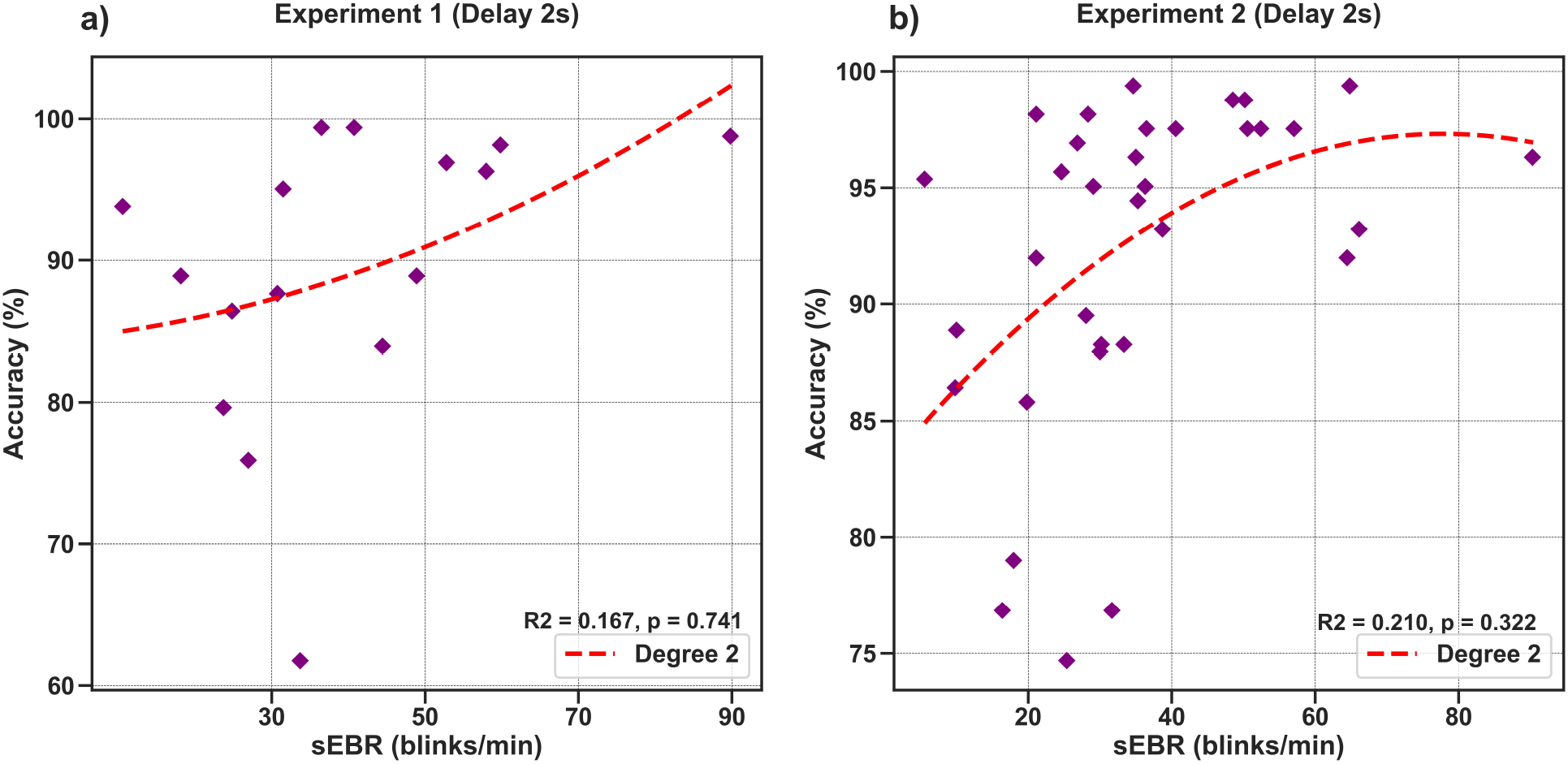
Polynomial regression model between task accuracy and sEBR during the first two seconds of the delay for Experiment 1 and Experiment 2. Regression plots show sEBR during the first two seconds of the Delay on the x-axis and task accuracy on the y-axis. (**A)** Polynomial regression model fitted on sEBR during the Delay and task accuracy in Experiment 1. (**B)** Polynomial regression model fitted on sEBR during the Delay and task accuracy in Experiment 2. Fitted red line represents polynomial regression model fit. The relationship between sEBR and WM performance appears to be non-linear and explains about 20% of the variance in Experiment 2 but does not reach significance. * p < .05, ** p < .01, *** p < .001.

### 3.4 Statistical Power

Post-hoc statistical power calculations using as the effect size the correlation between sEBR during the Delay period and task accuracy showed inadequate power for experiment 1 (N=16, 1-β error probability = 0.53, critical r=0.49). Power for experiment 2 was good (N=34, 1-β error probability = 0.87, critical r=0.33). Given that experiments 1 and 2 utilized different methods of quantifying blinks (camera-based vs. EOG) but a similar task design, the two sample sizes were combined with very good power obtained (N=50, 1-β error probability = 0.96, critical r=0.27).

## 4 Discussion

In the present study, we investigated the temporal fluctuations in sEBR across a WM paradigm and its relation to WM task accuracy in two experiments, inside and outside an MRI scanner, and using two methods of collecting sEBR. Using the same Sternberg working memory paradigm, we observed a strong positive relationship between sEBR and task performance only during the WM task delay in both experiments. We also found a significant difference in sEBR between task phases and a difference between Delay period sEBR and baseline sEBR.

Our first hypothesis was that phasic sEBR during the Delay period of the WM task would be positively correlated with task accuracy and that we would also observe a non-linear relationship where high and low sEBR would be predictive of low performance. We observed a strong positive correlation between sEBR during the Delay period in both Experiment 1 and Experiment 2 with task accuracy. However, only in Experiment 2 was this relationship significant. We believe that the lack of significance in Experiment 1 is due to the smaller sample size and thus lack of power, which our formal post-hoc power analyses confirmed. While the sample size in Experiment 2 was also small, we observed a similar correlation and higher power while recording sEBR using a different method (electrooculogram instead of camera-based eye-tracking hardware). The replication of a similar correlation between Delay period sEBR and WM performance across two separate experiments strengthens our findings. However, previous research has found that correlations begin to stabilize at even larger sample sizes(Schönbrodt and Perugini, 2013). Thus, future studies should aim to replicate these findings to investigate whether the observed effect stabilizes with even larger sample sizes.

If we interpret sEBR as an indirect measure of striatal DA activity, as other studies have postulated, we can speculate that higher sEBR during the WM delay was correlated with task accuracy due to DA regulating the maintenance and updating of representations in WM (Westbrook and Braver, 2016). The other results support this idea since no other task period was significantly correlated with task accuracy. Many studies that have investigated the relationship between sEBR and cognitive functions have used baseline levels of sEBR taken before or after tasks in their analysis (Tharp and Pickering, 2011;Zhang et al., 2015;Unsworth et al., 2019b). However, we show that while the WM task Delay period sEBR was correlated positively with task accuracy, baseline levels of sEBR were not. Our results highlight the importance of examining phasic and tonic sEBR when investigating the relationships between sEBR and other cognitive functions. The results also highlight that blinking may be an important component of working memory function, however, future studies, including within-subject analyses using larger number of trials, are needed to understand the role of blinking during WM maintenance. Additionally, future studies should investigate whether higher blink rates during the WM delay leads to a correct response. Since task difficulty was not controlled for in this study, participants’ task performance in both experiments was relatively high (see Tables 1 and 2). These limitations make our current dataset incapable of investigating these questions.

We also investigated the proposed “Inverted-U-shape” relationship between DA and WM performance by computing a polynomial regression model on sEBR during the delay and task accuracy (Cools and D’Esposito, 2011). Though the model showed a non-linear trend in Experiment 2, the model was not significant. We believe that failure to achieve non-linear model significance was due to lack of extreme (sub- and supraoptimal) sEBRs observed in the pool of participants, which are typically found in clinical populations (e.g., with Schizophrenia) (Adamson, 1995;Swarztrauber and Fujikawa, 1998). Future studies should investigate sEBR with healthy subjects and with subjects that have been observed to have extreme sEBR in order to have a wider variety of sEBRs and to better understand its connection with DA. Additionally, other methods of DA measures could be used to investigate DA during the delay period such as correlations with neuromelanin-sensitive MRI which can detect neuromelanin, a product of dopamine metabolism (Cassidy et al., 2019).

Our second hypothesis was that we would see significant differences in sEBR across the WM task as well as between sEBR during Rest and during the Delay period. We found sEBR to be the lowest during periods like Encoding and Probe in both Experiments, while sEBR during the Delay was the highest. Our results support previous findings which found task-related modulation of sEBR (Siegle et al., 2008;Oh et al., 2012). Prior work has found sEBR to be lower during tasks that require visual attention (Fukuda et al., 2005;Oh et al., 2012). This would explain the lower sEBR’s observed during the Encoding period when participants are encoding information into WM and during the Probe period where participants are retrieving information from WM. We also found that sEBR was the highest during the Delay period when participants were maintaining information in WM. This was also demonstrated in a different study which investigated sEBR during an A-not-B WM task where infants had to search for a hidden toy by making an eye movement to one of two locations (Bacher et al., 2017). Higher sEBR during the WM delay could be due to DA regulating the maintenance and updating of representations in WM (Westbrook and Braver, 2016), but this remains speculation until further studies directly measure dopaminergic activity during task performance. Our results further support this interpretation since Delay period sEBR was significantly higher than baseline sEBR during the Rest period. Lower sEBR during the Rest period could be explained since there is no need to update or maintain WM during this period.

To conclude, we investigated temporal changes of sEBR during different phases of a WM task and its relation to WM performance. We observed a significant positive correlation between sEBR and WM task performance during the Delay period, but not during the other phases of the task. Additionally, we found evidence for an association of sEBR during both stimulus encoding and WM probe retrieval, potential reflecting visual attention. To the best of our knowledge, this is the first study to investigate phasic and tonic sEBR during different phases of a WM task using complex visual scenes. Future studies should continue to investigate sEBRs in relation to direct measures of cortical (especially PFC) and subcortical dopamine and assess linear and non-linear relationships to task performance in healthy and clinical populations (e.g., Schizophrenia and Parkinson’s disease).

## Acknowledgments

Research reported in this publication was supported by the National Institute of Mental Health of the National Institutes of Health under Award Number R56MH116007 (T.M.E.).

## Author contributions

T.M.E designed the study. J.O, C.R, and B.G performed the research. J.O analyzed and interpreted the data, prepared the figures, and wrote the final manuscript. C.R, B.G, and T.M.E edited and reviewed the manuscript.

## Competing interests

The authors declare no competing interests.

## Notes

### Competing Interest Statement

The authors have declared no competing interest.

### Summary of Updates

This revision contains additional analyses and expanded discussion.

